# Microbiome functional gene pathways predict cognitive performance in older adults with Alzheimer’s disease

**DOI:** 10.1101/2025.03.06.641911

**Authors:** Abigail L. Zeamer, Yushuan Lai, Victoria Sanborn, Ethan Loew, Matthew Tracy, Cynthia Jo, Doyle V. Ward, Shakti K. Bhattarai, Johnathan Drake, Beth A. McCormick, Vanni Bucci, John P. Haran

**Author notes:** CORRESPONDING AUTHOR: John P. Haran.

## Abstract

Disturbances in the gut microbiome is increasing correlated with neurodegenerative disorders, including Alzheimer’s Disease. The microbiome may in fact influence disease pathology in AD by triggering or potentiating systemic and neuroinflammation, thereby driving disease pathology along the “microbiota-gut-brain-axis”. Currently, drivers of cognitive decline and symptomatic progression in AD remain unknown and understudied. Changes in gut microbiome composition may offer clues to potential systemic physiologic and neuropathologic changes that contribute to cognitive decline. Here, we recruited a cohort of 260 older adults (age 60+) living in the community and followed them over time, tracking objective measures of cognition, clinical information, and gut microbiomes. Subjects were classified as healthy controls or as having mild cognitive impairment based on cognitive performance. Those with a diagnosis of Alzheimer’s Diseases with confirmed using serum biomarkers. Using metagenomic sequencing, we found that relative species abundances correlated well with cognition status (MCI or AD). Furthermore, gene pathways analyses suggest certain microbial metabolic pathways to either be correlated with cognitive decline or maintaining cognitive function. Specifically, genes involved in the urea cycle or production of methionine and cysteine predicted worse cognitive performance. Our study suggests that gut microbiome composition may predict AD cognitive performance.

## INTRODUCTION

Alzheimer’s disease (AD) is a highly prevalent and progressive neurodegenerative disease and the most common cause of dementia in older adults with no known prevention or cure^1^. AD pathology is characterized by beta-amyloid plaques, tau-based aggregates that form neurofibrillary tangles (NFTs), neuroinflammation, and overall gross neuronal atrophy. In individuals with AD, symptoms often appear years after changes in the brain have begun^2–4^. In the early stages of typical AD, impairment in episodic memory and executive function of varying severity are prototypical. As the disease progresses and impacts additional parts of the brain, other cognitive domains show progressive impairment, including semantic memory, language and visuospatial domains^5^. In combination with symptoms, AD can be diagnosed and staged using a variety of imaging and blood-based biomarkers^3^. Despite extensive development and study, targeting beta-amyloid aggregates has shown minimal improvements in cognition or halting disease progression^6^. The minimal success of these therapies and a high disease burden warrant the exploration of other avenues of intervention.

Neuroinflammation plays a crucial role in managing the spread of injury and inflammatory molecules by addressing both the initial cause of injury and the resulting damaged tissue and cells^7^. Microglia, the resident innate immune cell of the brain, become increasingly activated with age and are activated by the binding of misfolded and aggregated peptides (DAMPs) or microbial-derived molecules (PAMPs) to pattern recognition receptors (PRRs) expressed on resting microglia^8,9^. The pro-inflammatory response initiated by PAMP/DAMP binding diverts the microglia from homeostatic function towards damage response pathways^10^. When activation is sustained by continued exposure to beta-amyloid or microbial products, excess pro-inflammatory cytokines such as IL-1 and IL-18 lead to neurodegeneration^11^. In the periphery, extracellular microbial products, including lipopolysaccharide (LPS) and peptidoglycan (PG), initiate an innate immune response through MD-2/TLR4 and related pathways to stimulate the release of IL-1β and IL-18.

The gut microbiome is one of the largest sources of microbial metabolites in the human body. Disruptions of microbiome composition and diversity (termed dysbiosis) are associated with multiple neurodegenerative conditions that also have a strong neuroinflammatory component, including Multiple Sclerosis, Parkinson’s Disease, and AD^12–14^. The microbiota-gut-brain (MGB) axis describes the bi-directional communication of the gut microbes and the central nervous system through cytokines, hormones, neuronal signals, and bacteria-derived molecules^15–17^. While the blood-brain barrier (BBB) prevents the free flow of microbial metabolites to the brain, age, and chronic inflammation, the BBB and gut barriers weaken, enabling dysregulation of the MGB axis and allowing microbes and microbial products to infiltrate the brain^18^.

The microbiota of individuals with AD has previously been described and characterized using a binary diagnosis of AD or not^14,19,20^. Broadly, Vogt and authors^14^ reported an increased abundance of *Bacteroidetes* and a lower abundance of Firmicutes and Bifidobacterium species in AD patients. In these studies, AD subjects are usually at similar stages of disease and are sampled at a single timepoint, thus failing to provide any perspective on the temporality of changes in microbiota and cognition. Outcomes that better represent AD are needed to dissect the relationship between AD and the microbiome, outcomes that better represent AD are needed. Instead of using a binary diagnosis of AD or not, measures of severity and cognitive staging can be analyzed to provide a better representation of disease.

Progress or stage of Alzheimer’s disease is commonly measured using mental status scales such as the Mini-Mental State Examination (MMSE) and clinical dementia rating (CDR)^21,22^. In this study we used the Alzheimer’s Disease Assessment Scale Cognitive subscale 13 item (ADAS-Cog-13) test as it is commonly used in clinical trials and longitudinal studies where it shows high sensitivity and ability to track AD specific progression^23^. The ADAS-Cog-13 is a global cognitive measure attained the combination of 13 tasks covering the cognitive domains of memory, language, praxis and executive functioning^24^.

Here we utilize the Gut-brain Alzheimer’s disease Inflammation and Neurocognitive Study (GAINS)^25^ to explore the microbiome of older individuals over time and in relationship to environment and cognitive function. Within this cohort, we found that gut microbiome composition correlated well with objective measures of cognition in both the MCI and AD groups. Analysis of gene pathways also point to a “functional” microbiome in which select microbial metabolites is correlated with cognitive decline.

## MATERIALS AND METHODS

### Study design and subject categorization

Older individuals (60 years or older) residing independently in Massachusetts were recruited to the Gut-brain Alzheimer’s Disease Inflammation and Neurocognitive Study (GAINS). During enrollment, demographics (i.e., age, sex, race, and education) and medical history were collected. At enrollment and each follow-up visit (occurring every 90 days), fecal samples, nutritional status (using the Mini-Nutritional Assessment^26^), frailty (measured using the Canadian Study of Health and Aging’s 7-point Clinical Frailty Scale (CFS)^27^, hospital exposure and medications were collected. We also performed a battery of cognitive assessments at each visit including the Alzheimer’s Disease Assessment Scale-Cognitive Subscale 13 (ADAS-Cog-13)^24^, the Clinical Dementia Rating Scale (CDR)^25^, and the Cognitive Battery (CB) of the National Institutes of Health (NIH) Toolbox for the assessment of neurological and behavioral function^28,29^.

Subjects were categorized into three groupings post-hoc: Alzheimer’s disease (AD), mildly cognitively impaired (MCI), and cognitively normal healthy controls (HC). For subjects to be labeled as having AD, they or their caregiver self-reported a diagnosis by a clinician. To confirm a diagnosis, research staff reviewed medical records for neurological assessments and neuroimaging. Subjects were classified as having MCI if the met all of the following criteria: Word List Delayed Recall score >1 SD above mean; 2) normal mental status: Mini-Mental State Examination score 25+; 3) normal daily functioning; 4) memory complaint: subjective or family response to standardized question; 5) not demented: CDR score <1 was used for subjects whose scores for greater than or equal to four for the word list delayed recall task of the ADAS-Cog-13^30^. Subjects without either a dementia diagnosis or impaired cognitive performance, were labeled as HC for healthy control.

### Cognitive scores and the derivation of cognitive domain specific z-scores

The ADAS-Cog-13 subscale was originally developed as an extension of the original 11-item ADAS-Cog, which added tasks to assess attention and concentration, planning and executive function, verbal and nonverbal memory, and praxis^24,31^. Scores range from 0 to 85, with higher scores indicative of higher cognitive impairment.

To examine whether performance in the two early impaired AD-specific cognitive domains of memory and executive function is associated with microbiome composition, we created composite z-scores to ease interpretation and comparison across domains. For our memory z-score, we calculated the sum at each sample point of the word recall score, word recognition score, orientation score, remembering word recognition score, and delayed recall score. The mean and standard deviation of the sum score of all subjects at each visit time were calculated. A z-score for each visit was calculated by subtracting the mean memory sum score of the visit from the memory sum score of a subject and dividing it by the standard deviation of that visit. An executive function z-score was found by first summing the executive function maze time score, maze number of mistake score, dimensional card sort task (NIH toolbox), and the pattern comparison processing speed test score (NIH toolbox). The mean and standard deviation of each group of tests were calculated for each visit group and used to calculate a visit and sample-specific z-score. For both measures, a higher value is indicative of greater cognitive impairment.

### Sample handling and DNA sequencing

Stool samples were collected at home and mailed to the lab. All stool samples were collected as part of the participant’s normal bowel habits from defecation using the Fisherbrand™ Commode Specimen Collection System and sample placed into the OMNIgene•GUT collection kits (DNAgenotek, catalog no. OMR-200) which stabilizes the sample at room temperature for shipment via mail service.

Samples were stored at -80°C until DNA extraction and sequencing were performed. DNA was extracted from samples using the QIAGEN DNAeasy PowerSoil kits (QIAGEN, catalog no. 47016). Sequencing libraries were constructed using the Nextera XT DNA Library Prep Kit. Libraries were sequenced on a NextSeq 500 or NextSeq 1000 system with 2×150 nucleotide paired-end reads.

Remaining reads were trimmed and filtered for host contamination using the KneadData pipeline (https://github.com/biobakery/kneaddata). Metagenomic reads were profiled for microbial abundance, metabolic pathways and other functional measures using MetaPhlAn4 and HUMAnN3 databases and tools^32,33^.

### Microbiome analysis and modeling

#### Clinical covariate contribution to changing microbiome composition using MERF

As performed in^34^, mixed-effect random forest (MERF) regression modeling was used to determine the contribution of clinical covariates to microbial composition. For every microbe, the relative abundance was predicted as a function of clinical covariates and study ID as a random effect. Permutated variable importance was used to determine which clinical covariates were significantly predictive of microbiome composition (p-value cut-off < 0.05).

#### Linear mixed-effect modeling for diversity and cognitive status

To identify the influence of sex, age, education, antibiotic exposure, time since enrollment, and cognitive status on microbiome diversity (Inverse Simpson), we implemented a linear mixed-effects model with the following formula:

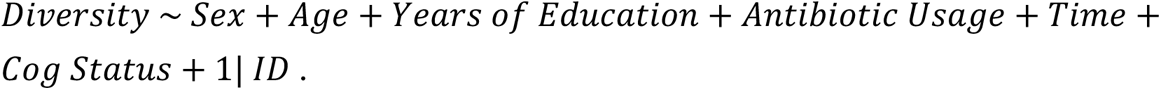

Linear mixed-effect models were implemented with the nlme R package. The random effect 1|*ID* was included to account for individual variation. The alpha diversity index, inverse Simpson, was calculated using the phyloseq R package. Sex represents what the subject identifies as at enrollment. In this study, all participants identified with what they were assigned at birth. Antibiotic usage showed whether the subject had used antibiotics in the 6 months preceding sampling and time represented days since enrollment.

To assess the stability of cognitive scores with time we again implemented linear mixed-effect models. In the entire GAINS cohort, the following formula was used:

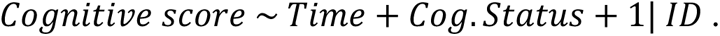

#### MERF modeling of ADAS-Cog-13 and cognitive domain z-scores

To find associations between cognitive scores and microbiome features, we developed a MERF regression pipeline. The first step of this pipeline was to split the longitudinal data into a train and test set by randomly leaving out one sample from every subject from the training set to be used to build the test set. Feature selection was first performed on the training data using the Boruta algorithm^35^. Selected features were then used to build a MERF model^36^. The performance of the models are presented as actual versus predicted graphs, where the predicted data is from the test data set for each seed. The procedure was repeated ten times, corresponding to ten different random seeds. A final version of the model was generated using all Boruta-selected features across the ten seeds and all subject samples. Permutated variable importance was then run using the final model^37^. All analysis and visualizations were performed using R.

## RESULTS

### Characteristics of the GAINS cohort participants

Participants were recruited to the GAINS from community centers and physicians’ offices across central Massachusetts. Over four years, 223 participants provided fecal samples at least once, with an average of 3.8 (SD 2.4) samples collected per participant for 856 total samples. The average age of participants was 71.5 years (SD 7.5) and 63.7% were female (**Table 1**). Seventeen and a half percent of participants were exposed to antibiotics in the 6 months preceding sampling (**Table 1**). Polypharmacy was prevalent in 34.1% of participants, with 46.6% of participants taking statins (**Table 1 and Supplemental Table 1**).

**Table 1:** GAINS patient demographics and clinical scores.

| **Elder**<br>**Characteristic<sup>a</sup>** | **Cog.**<br>**Normal**<br>**(n=158)** | **MCI**<br>**(n=40)** | **AD**<br>**(n=25)** | **P-value** |
| --- | --- | --- | --- | --- |
| Age (Mean [SD])<br>(years) | 70.2<br>(7.46) | 75.1<br>(7.22) | 73.9<br>(5.75) | <0.001 |
| **Sex** |  |  |  |  |
| Female | 109<br>(69.0%) | 21<br>(52.5%) | 12<br>(48.0%) | 0.0805 |
| Male | 49<br>(31.0%) | 19<br>(47.5%) | 13<br>(52.0%) |  |
| **Polypharmacy** |  |  |  |  |
| Less than 5<br>medications | 115<br>(72.8%) | 24<br>(60.0%) | 8 (32.0%) | <0.001 |
| 5 or more<br>medications | 43<br>(27.2%) | 16<br>(40.0%) | 17<br>(68.0%) |  |
| Antibiotics (past<br>6mon) | 28<br>(17.7%) | 9 (22.5%) | 2 (8.0%) | 0.475 |
| Hospital<br>Exposure | 13 (8.2%) | 4 (10.0%) | 1 (4.0%) | 0.753 |
| **Clinical Scores** |  |  |  |  |
| CFS (Mean [SD]) | 2.07<br>(1.02) | 2.54<br>(0.942) | 3.52<br>(1.53) | <0.001 |
| MIS (Mean [SD]) | 1.21<br>(0.470) | 1.28<br>(0.510) | 1.68<br>(0.627) | <0.001 |
| ADAS-Cog-13<br>(Mean [SD]) | 9.01<br>(4.42) | 21.0<br>(12.8) | 32.3<br>(20.1) | <0.001 |
| <sup>a</sup>Data are presented as the number (%), unless otherwise specified. |  |  |  |  |

Based on information collected at the first clinic visit, participants were segregated into one of three cognitive status groups: Cognitively normal healthy controls (HC), mild cognitively impaired (MCI), and Alzheimer’s Disease (AD) (see Methods section for further explanation). Age and occurrence of polypharmacy differed significantly by status, with the MCI group being older than the others and the AD group having a larger percentage of members taking more than five medications daily (Table 1). Within the GAINS cohort, alpha diversity was not significantly different across different cognitive status (**Supplemental Figure 1**). However, antibiotic usage in the past 6 months was significantly associated with lower diversity (Supplemental Table 2).

### Microbial composition is informed by demographics and clinical factors in the GAINS cohort

To examine associations between microbial abundance and clinical covariates in the GAINS cohort, we ran mixed-effect random forest (MERF) modeling to predict each species abundance as a function of demographics and clinical covariates. A mix of demographics, medications, and environmental factors was significantly predictive of microbial abundance across the 311 microbe-specific models (FDR adjusted p-value < 0.05; **Figure 1**). While age was the most frequently selected feature (appearing in 57.2% of models), medication accounted for 54.79% of the significant predictors. Specifically, psychoactive and antihypertensive medications were most prevalently associated with microbiome abundance.

**Figure 1:**
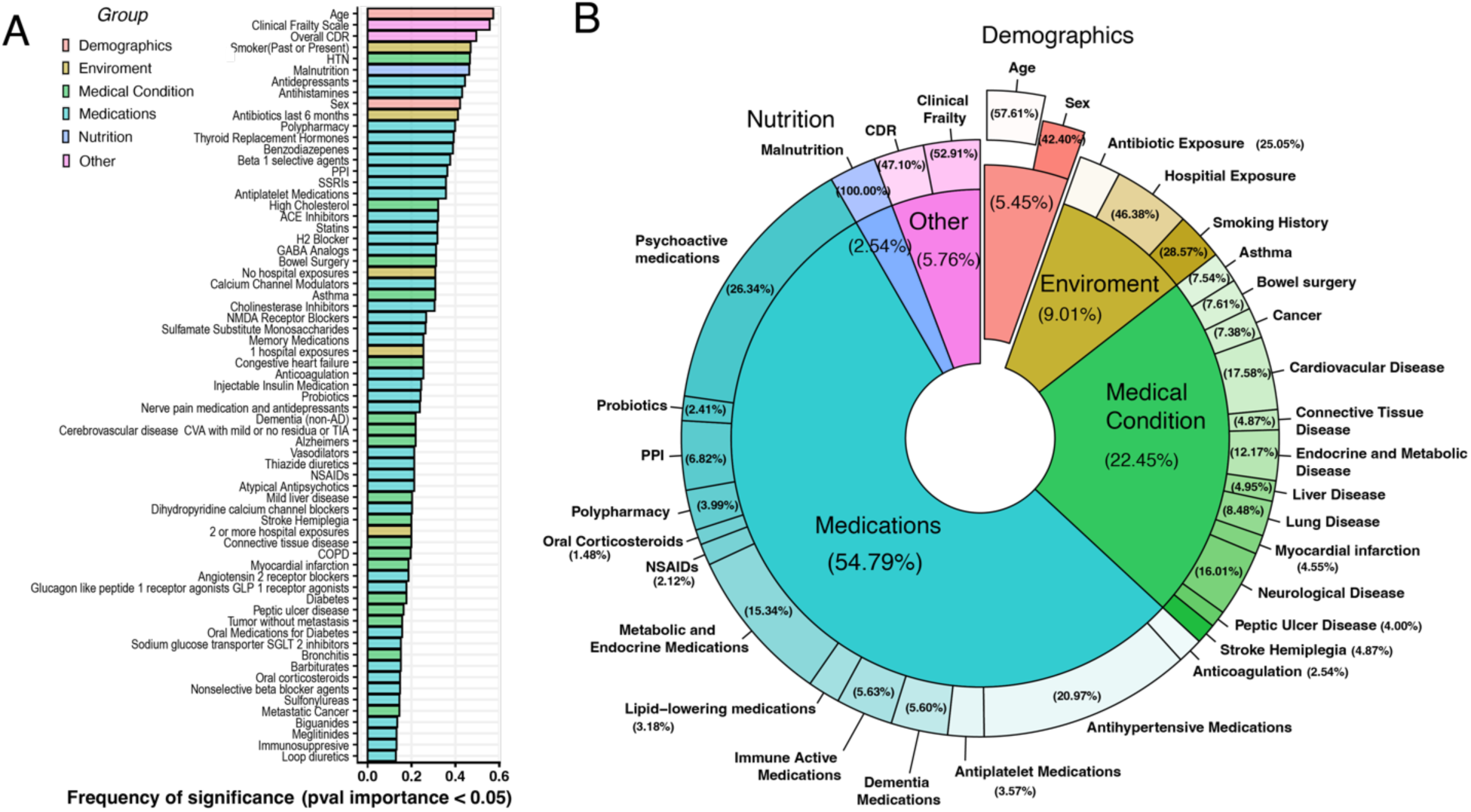
Clinical and demographic variables that predict microbiome composition in community dwelling individuals. MERF modeling results using all demographics and clinical variables (including each medication taken and clinical scoring of frailty, nutrition, and comorbidities) to predict each species abundance from repeated fecal samples. (A) Frequency-based ranking of predictors from MERF models. Significance was determined by running permutated variable importance analysis and using an FDR adjusted p-value of 0.05. (B) Pie chart to illustrate the distribution of the clinical factors found by the modeling to be significantly associated with the GAINS microbiome. Drugs were grouped by treatment class.

### Cognitive measures show stability in the AD participants

In the GAINS cohort, a battery of cognitive testing was performed at each visit and provided us with three measures of interest: ADAS-Cog-13, memory z-score, and executive function (EF) z-score. The ADAS-Cog-13 is a global measure of cognitive function with a specific emphasis on domains implicated in AD progression. Memory and EF z-scores are specific to each respective cognitive domain and are derived from tasks originating from the ADAS-Cog-13 and NIH toolbox (source and derivation are further explained in the *Methods* section). Linear mixed-effect models where cognitive scores were modeled as a function of sample day, cognitive status, and study ID found that ADAS-Cog-13 scores and memory z-scores, but not EF z-score, reduced with time (**Table 2**). In the MCI and HC subgroups, ADAS-Cog scores varied significantly with time (**Table 3**). Memory z-score varied significantly in the MCI cohort with time (**Table 3**). In the AD group, none of the cognitive scores varied significantly with time, suggesting greater stability in the AD cohort compared with the HC or MCI cohorts (**Table 3**).

**TABLE 2:**
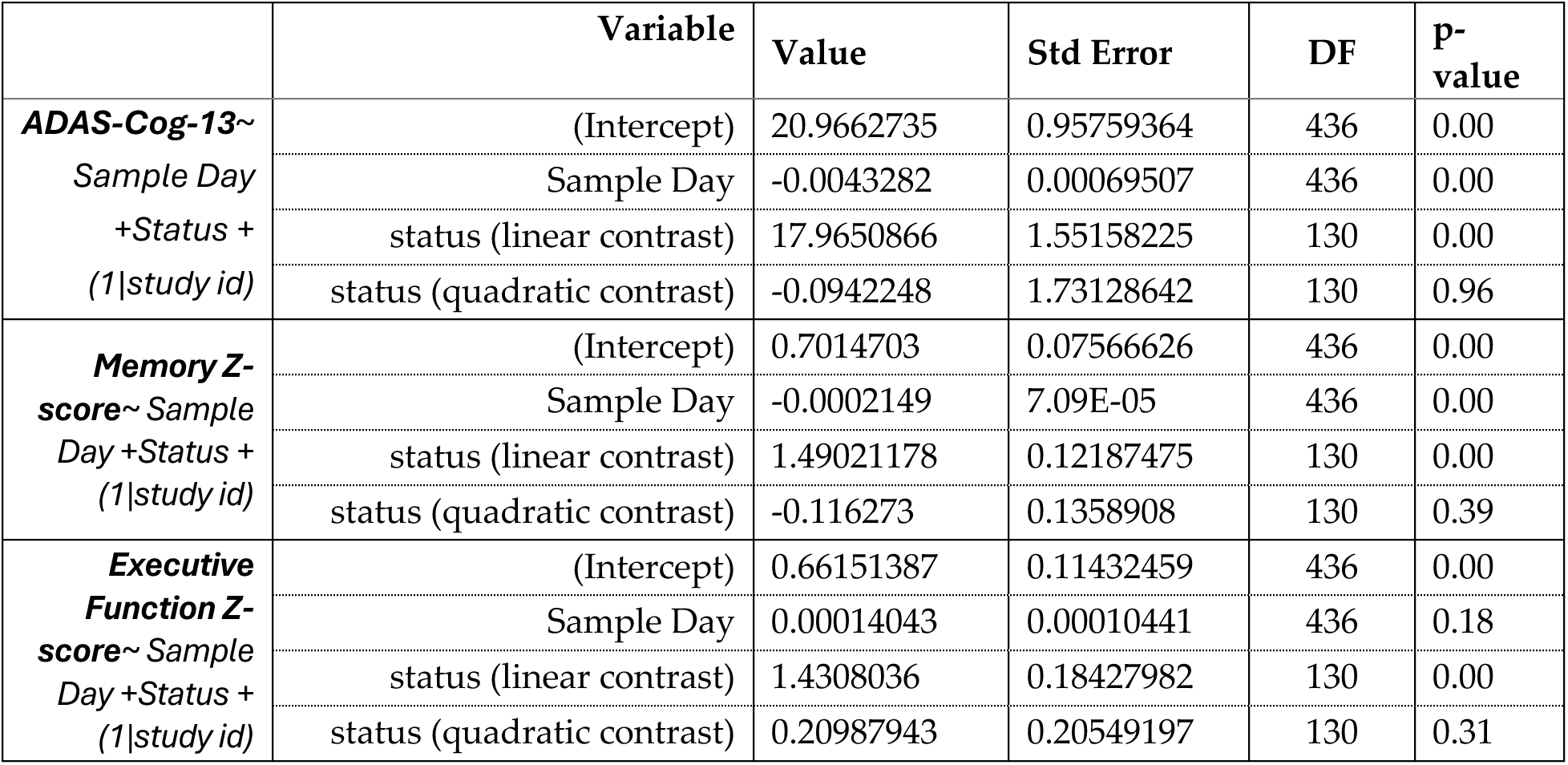
LLM results of cognitive measures in GAINS.

|  | Variable | Value | Std Error | DF | P-value |
| --- | --- | --- | --- | --- | --- |
| **ADAS-Cog-13~**<br>*Sample Day*<br>**+Status +**<br>*(1\|study id)* | (Intercept) | 20.9662735 | 0.95759364 | 436 | 0.00 |
|  | Sample Day | -0.0043282 | 0.00069507 | 436 | 0.00 |
|  | status (linear contrast) | 17.9650866 | 1.55158225 | 130 | 0.00 |
|  | status (quadratic contrast) | -0.0942248 | 1.73128642 | 130 | 0.96 |
| **Memory Z-score~**<br>*Sample Day*<br>**+Status +**<br>*(1\|study id)* | (Intercept) | 0.7014703 | 0.07566626 | 436 | 0.00 |
|  | Sample Day | -0.0002149 | 7.09E-05 | 436 | 0.00 |
|  | status (linear contrast) | 1.49021178 | 0.12187475 | 130 | 0.00 |
|  | status (quadratic contrast) | -0.116273 | 0.1358908 | 130 | 0.39 |
| **Executive Function Z-score~**<br>*Sample Day*<br>**+Status +**<br>*(1\|study id)* | (Intercept) | 0.66151387 | 0.11432459 | 436 | 0.00 |
|  | Sample Day | 0.00014043 | 0.00010441 | 436 | 0.18 |
|  | status (linear contrast) | 1.4308036 | 0.18427982 | 130 | 0.00 |
|  | status (quadratic contrast) | 0.20987943 | 0.20549197 | 130 | 0.31 |

**TABLE 3:** LLM results of cognitive measures predicted by cognitive status.

|  | Group | Variable | Value | Std Error | DF | p-value |
| --- | --- | --- | --- | --- | --- | --- |
| **ADAS-Cog-13**<br>*~ Sample Day + (1\|study id)* | HC | (Intercept) | 8.283773 | 0.37480035 | 325 | 1E-66 |
|  |  | Sample Day | -0.0045605 | 0.00068083 | 325 | 9E-11 |
|  | MCI | (Intercept) | 21.9278189 | 2.81897912 | 61 | 0.00000 |
|  |  | Sample Day | -0.0086379 | 0.00215153 | 61 | 0.00017 |
|  | AD | (Intercept) | 32.2640195 | 4.61882516 | 48 | 0.00 |
|  |  | Sample Day | 0.00350906 | 0.00296073 | 48 | 0.24 |
| **Memory Z-score**<br>*~ Sample Day + (1\|study id)* | HC | (Intercept) | -0.4204712 | 0.03619103 | 325 | 0.00 |
|  |  | Sample Day | -8.61E-05 | 7.27E-05 | 325 | 0.24 |
|  | MCI | (Intercept) | 0.90001334 | 0.21765386 | 61 | 0.0001 |
|  |  | Sample Day | -0.0007137 | 0.00021877 | 61 | 0.0018 |
|  | AD | (Intercept) | 1.74257229 | 0.34399324 | 48 | 0.00 |
|  |  | Sample Day | -0.0003959 | 0.00028079 | 48 | 0.17 |
| **Executive Fun. Z-score**<br>*~ Sample Day + (1\|study id)* | HC | (Intercept) | -0.259374 | 0.057333 | 325 | 0.000009 |
|  |  | Sample Day | 0.000107 | 0.000096 | 325 | 0.26 |
|  | MCI | (Intercept) | 0.468305 | 0.284264 | 61 | 0.10 |
|  |  | Sample Day | 0.000375 | 0.000273 | 61 | 0.17 |
|  | AD | (Intercept) | 1.746795 | 0.556368 | 48 | 0.003 |
|  |  | Sample Day | 0.000206 | 0.000586 | 48 | 0.73 |

### Demographics and clinical covariates correlate with cognitive measures

Performance on cognitive tests can be influenced by a myriad, of factors including age and years of education^38^. In the HC and MCI groups of participants, age positively correlated with our three cognitive measures (**Figure 2**). In the AD cohort, age was negatively correlated with all scores, suggesting that younger participants are more severely impaired than those who are older. A negative correlation was also found between years of education and cognitive score in the HC cohort but not the MCI or AD cohorts. Frailty positively correlated with all scores in the AD cohort; in the MCI and HC cohorts’ frailty negatively correlated with ADAS-Cog-13 and EF z-scores. Malnutrition negatively correlated with ADAS-Cog-13 and EF z-scores in the HC cohort and EF z-scores in the MCI cohort. Altogether our analysis suggests that in the MCI and HC cohorts, health and education measures are more informative of cognitive scores than in AD cohorts. In the AD cohort, younger age and increased frailty correlated with cognitive scores.

**Figure 2:**
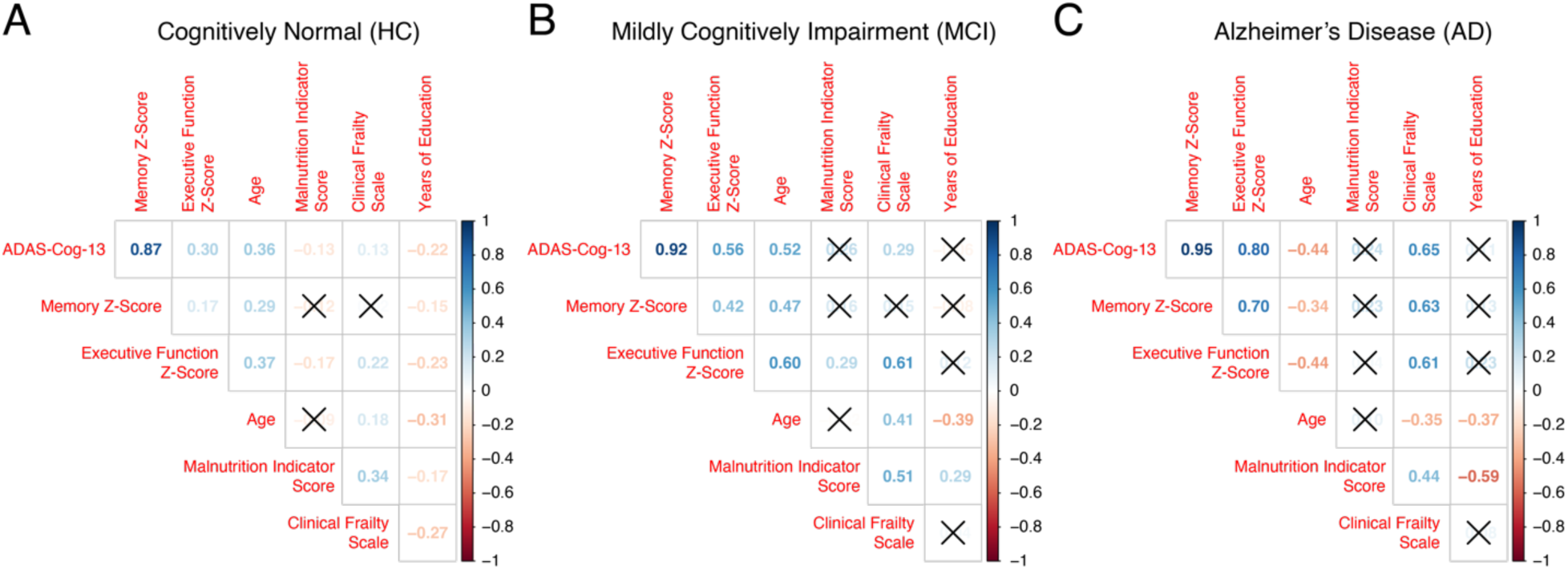
Spearmen correlation coefficient between each cognitive outcome (ADAS-Cog-13, Memory Z-score and Executive Function Z-scores) and relevant clinical covariates in (A) Cognitively normal, (B) Mildly cognitively impaired and (C) AD diagnosed participants.

### Depletion of health-associated microbes is informative of worse ADAS-Cog-13 scores only in individuals with MCI and AD

We then sought to identify microbiome features possibly associated with ADAS-Cog-13 scores. As above, we built MERF regression models for each subject subcategory. Models predicting ADAS-Cog-13 scores in the HC subject subset produced marginally informative results, as evident by the correlation between actual and predicted ADAS-Cog-13 for the species abundance-based model, 0.64 ± 0.05 (**Figure 3A, Supplemental Table 3**). Oppositely, models trained on MCI and AD subsets demonstrated strong predictive performance with correlations coefficients respectively of 0.87 ± 0.05 and 0.94 ± 0.03 (**Figure 3B,C and Supplemental Table 3**). This may be indicative that people with MCI or diagnosed with AD may have a microbiome composition that is more reflective of their cognitive performance compared to HC individuals. Further, the ADAS-Cog-13 is powered to identify differences in mild AD cases and thus focuses on AD specific cognitive domains. The test is thus not powered to assess general cognition in HC people.

**Figure 3:**
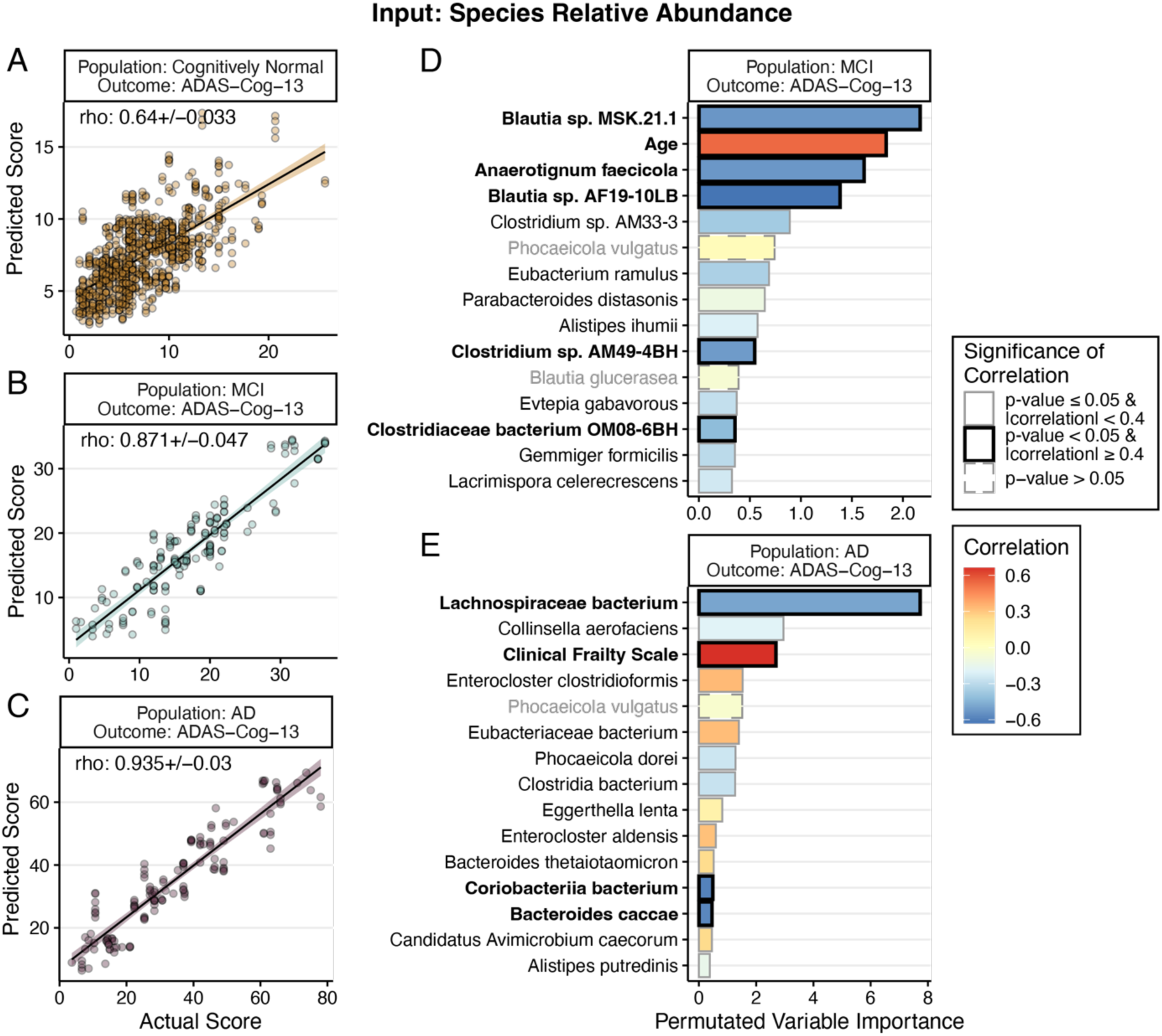
Mixed-Effect Random Forest (MERF) regression models using species abundance and clinical covariates to predict ADAS-Cog-13 scores. Actual versus predicted ADAS-Cog-13 values for cognitively normal (A), MCI (B) and AD patient populations (C). Permutated variable importance for the final model for the MCI (D) and AD (E) populations. Bar color (D and E) corresponds to the correlation coefficient of each variable with ADAS-Cog-13 score with red indicating positive correlation and blue indicating negative correlation. Significance of correlation is indicated by either a dashed (where p-value > 0.05) or a solid outline (p-value <= 0.5). Magnitude of correlation is indicated by color of the outline with gray indicating correlation below an absolute value of 0.4 and black indicating the correlation coefficient is great or equal to 0.4.

As the MCI and AD patient’s subsets cognitive performance-microbiome model displayed strong predictive power, we sought to determine the most informative microbial species of the predicted signal. Inspecting permutated importance resulting from the models we found that the leading microbial predictors of ADAS-Cog-13 scores in the MCI cohort included two *Blautia spps.* (a known health-associated Clostridium Cluster IV/XIVa SCFA-producing bacterium (Liu et al., 2021)) (ranked 1^st^ and 4^th^) and *Anaerotignum faecicola* (ranked 3^rd^), all of which showed significantly lower abundance correlating with worse/higher ADAS-Cog-13 scores (**Figure 3D and Supplemental Figure 2A**). Also, in the MCI cohort, increased age correlated with higher ADAS-Cog-13 scores and was identified as a top predictor (ranked 2^nd^), suggesting general age-related cognitive decline may be occurring in our MCI cohort.

Lower abundance of *Lachnospiraceae bacterium* (also a Clostridium cluster XIVa member)*, Coriobacteriia bacterium,* and *Bacteroides caccae* were found to be the top predictors of worse ADAS-Cog-13 scores in the AD population and were negatively correlated with score (**Figure 3E and Supplemental Figure 2B**). Corroborating the finding that frailty strongly correlates with ADAS-Cog-13 score (**as in Figure 2C**), the RF model also identified higher frailty as a stronger predictor of higher ADAS-Cog-13 scores in AD patients.

### Relative species abundance predicts performance in cognitive subdomains

The ADAS-Cog-13 score summarizes performance across cognitive domains, including memory, language, executive function, attention, and praxis. As the changes in cognitive function in AD are distinct^5^, we sought to determine if we could disentangle the predictive ability of microbial features seen for ADAS-Cog-13 to the contribution to a specific cognitive domain. To this end, we calculated composite z-scores for memory and executive function using components of the ADAS-Cog-13 test and the NIH toolbox (see *Methods* for further explanation of derivation). As memory and executive function represent domains profoundly affected in AD during the early clinical stages, we focused on testing involving these domains and calculated a memory z-score and executive function (EF) z-score. Like the ADAS-Cog-13 score, higher memory and EF z-scores values indicate greater impairment in the given domain or worse performance.

MERF models trained to predict memory z-scores from species abundance predicted scores that tightly correlated with the actual scores for both the MCI (correlation = 0.815 ± 0.07) and AD (correlation = 0.929 ± 0.01) populations (**Figure 4A**). Predictions of EF z-scores using species abundance were also well correlated with actual scores in both the MCI (0.838 ± 0.04) and AD populations (0.84 ± 0.04) (Figure 4D). Predictions in the cognitively normal population were again subpar and thus not further examined (**Figure 4A**, **4D**).

**Figure 4:**
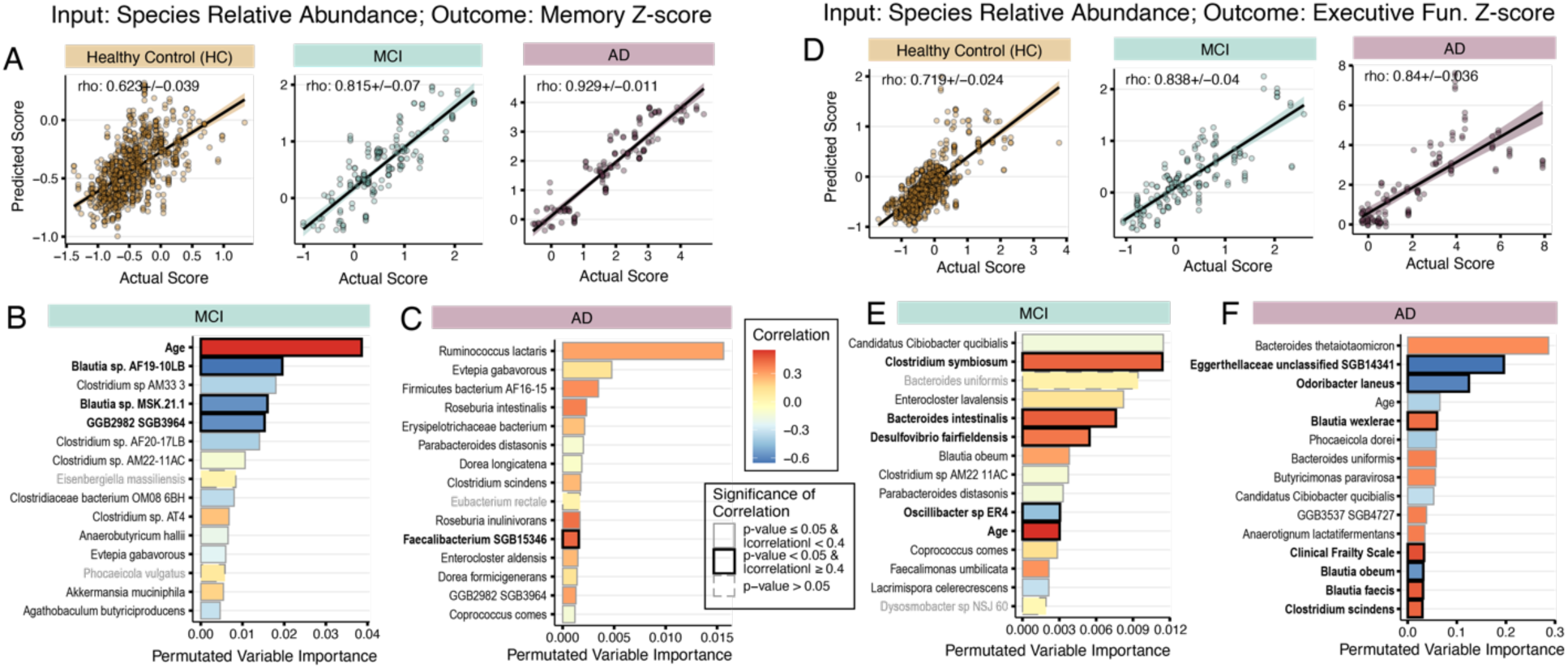
MERF regression models using species abundance and clinical covariates to predict memory z-scores and executive function z-scores. Actual versus predicted memory z-score values (A) for cognitively normal healthy controls (HC), mild cognitive impairment (MCI), and Alzheimer’s Disease (AD) patient populations. Permutated variable importance for the final model for the MCI (B) and AD (C) population. Actual versus predicted executive function z-score values (D) for HC, MCI, and AD patient populations. Permutated variable importance for the final model for the MCI (E) and AD (F) populations. Bar color (B, C, E, F) corresponds to the correlation coefficient of each variable with memory z-score were red indicates positive correlation and blue indicates negative correlation. Significance of correlation is indicated by either a dashed (where p-value > 0.05) or a solid outline (p-value <= 0.5). Magnitude of correlation is indicated by color of the outline with gray indicating correlation below an absolute value of 0.4 and black indicating the correlation coefficient is great or equal to 0.4.

Age was selected as a top predictor of both memory z-score and EF z-score in the MCI population (**Figure 4B**, **4E**). Like the ADAS-Cog-13 score, increased age significantly correlated with worse memory and EF z-scores, suggesting that some impairment may result from age-related cognitive impairment in the MCI population. To this end, the MCI cohort was significantly older (75.1 ± 7.22 years; p-value < 0.001) than both HC (70.2 ± 7.46 years) and AD cohorts (73.9 ± 5.75 years). In total, seven of the top fifteen predictors of memory z-score were selected by the ADAS-Cog-13 model. Six of the seven predictors significantly correlated with outcome including two *Blautia* species and two species belonging to the *Clostridiaceae* family which all showed decreased abundance correlating with higher scores (**Figure 5A**). *Phocaeicola vulgatus* and *Evtepia gabavorous* were identified as top predictors for the ADAS-Cog-13 and memory z-score but did not show significant or strong correlation with outcomes (correlation coefficient < |0.4|).

**Figure 5:**
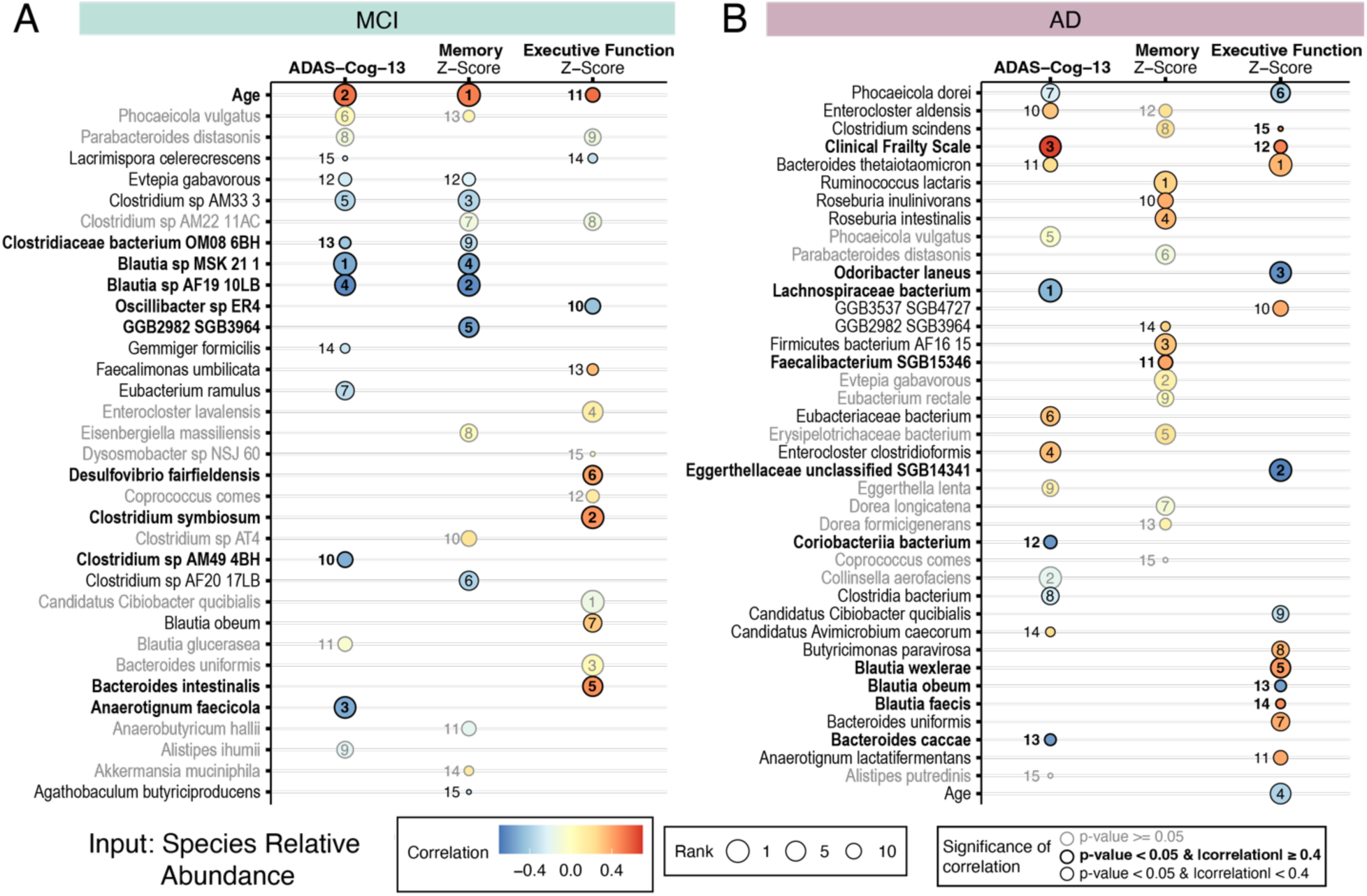
Comparison of rank and correlation of top 15 permutated variables from species abundance based MERF models in each cognitive outcome. Models trained with the MCI subset (A) and AD subsets (B). Gray coloring of microbial name, circle and number indicate that the correlation is not significant (p-value >= 0.05).

Compared to the memory z-score top predictors, fewer of the top predictors of EF z-score overlapped with the other cognitive models in the MCI population (**Figure 5A**). Only age was selected by both ADAS-Cog-13 and EF Z-score (**Supplemental Figure 2A and Supplemental Figure 4A**). Decreased abundance of a potential beneficial cholesterol metabolizing bacteria, *Oscillibacter* correlated with worse EF z-scores in the MCI cohort (Li et al., 2024). Increased abundance of *Clostridium symbiosum, Bacteroides intestinalis,* and *Desulfovibrio fairfieldensis* positively correlated with worse EF z-scores.

In the AD cohort model, the only species selected by the MERF model that also strongly correlated (correlation coefficient ≥0.4) is an uncultured *Faecalibacterium* species that showed increased abundance correlating with worse memory z-score (**Figure 4C**). *Enterocloster aldensis* (ranked 12^th^), which was also identified as important in predicting ADAS-Cog-13 did not show significant correlation with memory z-score (**Supplemental Figure 3B**).

Lower abundance of *Eggerthellaceae sp*. and *Blautia obeum* were identified as important predictors and correlated significantly with worse EF z-score in the AD group (**Figure 4F**). Conversely, higher abundance of *Blautia wexlerae* and *Blautia faecis* correlated with higher EF z-scores, suggesting a strain-specific association with cognition. Worse EF z-scores also correlated with a lower abundance of two important Bacteroides species, *Odoribacter laneus* and *Phocaeicola dorei* (**Supplemental Figure 4B**). Unlike in the MCI population, younger age positively correlated with worse EF z-score, suggesting that participants with a younger age of onset may be more severely impaired in the executive functioning domain than those with older ages of onset. A higher abundance of *Clostridium scindens* was identified as important in predicting worse performance (higher scores) for both EF and memory z-scores in the AD cohort. However, it was not significant for memory z-score (**Figure 5**).

### Enrichment of nitrogen-metabolizing and methionine-related pathways is informative of cognitive performance in AD participants

While informative of outcome, microbial abundance alone fails to provide insight into the functionality of the microbial community. Metabolic pathways enable a more complete view of the functional microbiome, or the function of genes found in fecal samples that produce metabolites in the gut and interact and influence the host. MERF models were run to predict ADAS-Cog-13, memory z-scores, and EF z-scores using basic clinical covariates and metabolic pathway relative abundance in AD participants. The predicted values tightly correlated with actual values for all three outcomes (ADAS-Cog-13 correlation = 0.935 ± 0.02; Memory z-score correlation = 0.944 ± 0.01; EF z-score correlation = 0.88 ± 0.03) (**Figure 6 A**).

**Figure 6:**
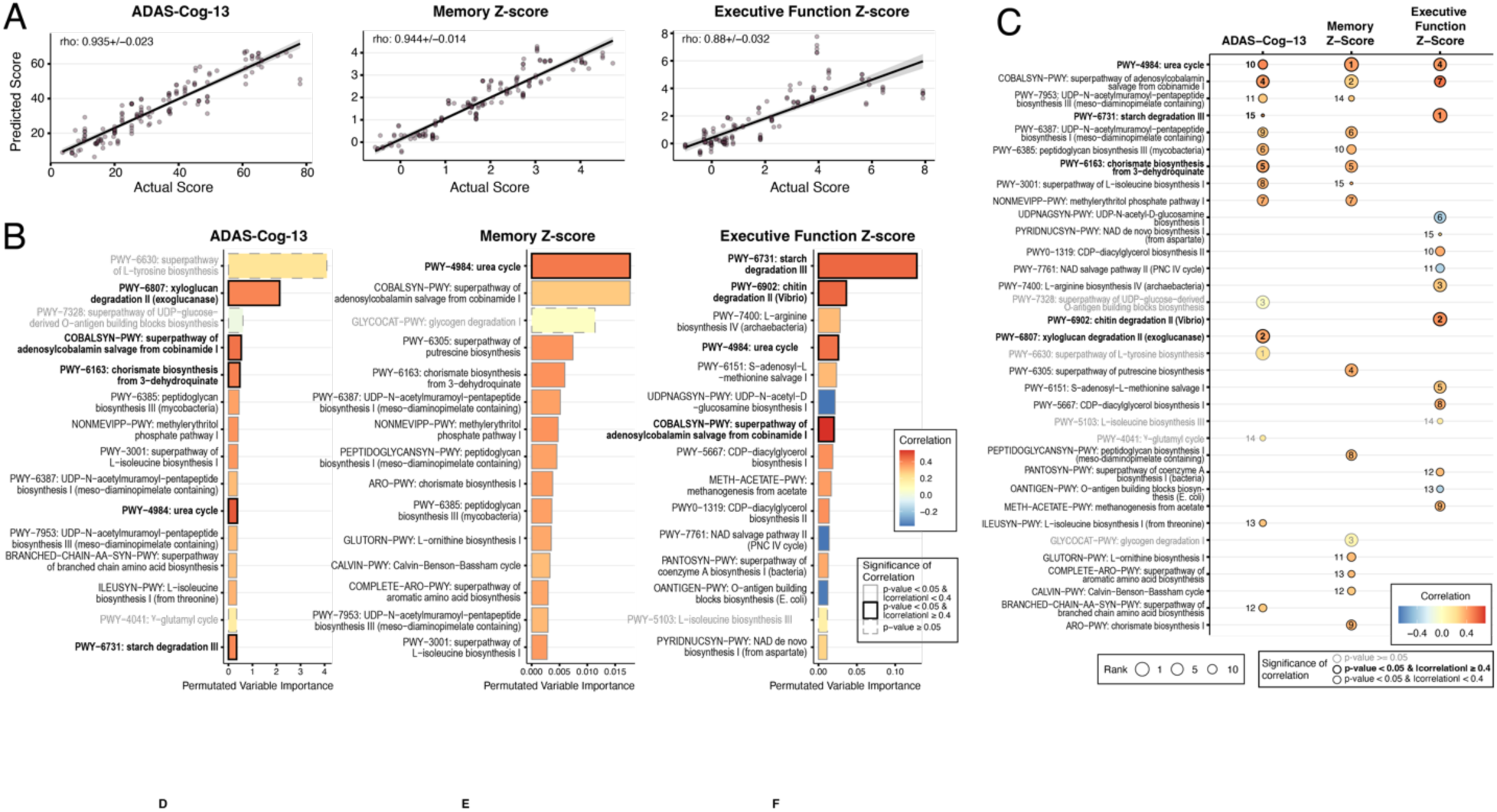
Pathway abundance analyses in AD population. Comparison of rank and correlation of top 15 permutated variables from pathway abundance based MERF models in each cognitive outcome. Models (A) trained with the AD subset to predict ADAS-Cog-13, memory z-score, and executive function z-score. Top 15 predictors (B) from ADAS-Cog-13, memory z-score, and executive function z-score colored for correlation coefficient of each pathway with the identified outcome. Significance of correlation is indicated by either a dashed (where p-value > 0.05) or a solid outline (p-value <= 0.5). Magnitude of correlation is indicated by color of the outline with gray indicating correlation below an absolute value of 0.4 and black indicating the correlation coefficient is great or equal to 0.4. (C) Comparison of rank and correlation of top 15 permutated variables from pathway abundance based MERF models in each cognitive outcome for the AD population. Gray coloring of microbial name, circle and number indicate that the correlation is not significant (p-value >= 0.05).

Across all three outcome models, ADAS-Cog-13 (ranked 10^th^), Memory z-score (ranked 1^st^), and EF z-score (ranked 4^th^), the urea cycle (PWY-4984) was selected as important with a higher abundance of the pathway correlating with worse performance in each cognitive outcome (**Figure 6 B**). While no other pathways directly involved in nitrogen assimilation were identified as important, two pathways involved in polyamine were important in the memory z-score model. Putrescine (PWY-6305) and ornithine (GLUTORN-PWY) biosynthesis specifically were selected as correlating with worse memory z-score but did not pass the correlation coefficient cut-off used of 0.4 (ranked 4^th^ and 11^th,^ respectively) (**Supplemental Figure 5B**).

A pathway for vitamin B12 cofactor salvaging (COBALSYN-PWY) was selected as important for all three outcomes of interest. A higher abundance of the salvaging pathway positively correlated with worse ADAS-Cog-13 scores (ranked 4^th^) and EF z-scores (ranked 7^th^). Vitamin B12 cofactor (adenosylcobalamin), the active form of vitamin B12 in the body, is a cofactor for highly conserved enzymes involved in amino acid synthesis and the breakdown of fatty and amino acids (Rowley and Kendall, 2019). In addition to vitamin B12 cofactor salvaging, three pathways involved in enzyme cofactor biosynthesis were important for predicting EF Z-score but did not meet the correlation cut-off (PYRIDNUCSYN -PWY, ranked 15^th^; PWY-7761, ranked 11^th^; PWY-6151, ranked 8^th^). A lower abundance of the NAD salvaging pathway (PWY-7761) correlated with worse EF z-scores, contrasting the trending towards a positive correlation of *de novo* NAD pathway (PYRIDNUCSYN-PWY) with EF z-score (**Supplemental Figure 5C**).

### Microbial genes involved in methionine and cysteine metabolism are informative of cognitive outcomes in AD participants

MetaCyc pathways, as used above, are well-curated and highly interpretable; however, understanding the interconnections between pathways and their relationships to specific outcomes often require complex, manual annotation. In contrast, the Kyoto Encyclopedia of Genes and Genomes (KEGG) ontology offers a broader coverage of pathways and functions that can enable a more holistic view of the microbiota. KEGG ontology (KO) terms represent functional orthologs that can be linked to a variety of larger pathways and groups. To identify specific gene functions associated with cognitive outcome in the AD subjects, we predicted our three outcomes using MERF models trained with KO terms encoded by the microbiome and basic clinical covariates Actual and predicted ADAS-Cog-13 (correlation=0.943 ± 0.02), memory z-score (correlation=0.949 ± 0.02), and EF z-score (correlation=0.87 ± 0.06) were tightly correlated, indicating high performance by the resulting models (**Figure 7A-C**). The top fifteen predictors of each outcome were combined, and enrichment analysis on the functionality of the selected KO terms was performed. Genes relating to cysteine and methionine metabolism were significantly enriched in the predictors, along with two genes relating to bacterial secretion system and protein export genes (**Figure 7G**). Enrichment of these two pathways suggests that individually or in combination, bacterial exported products, including methionine and cysteine, are predictive of cognitive outcomes in our AD cohort.

**Figure 7:**
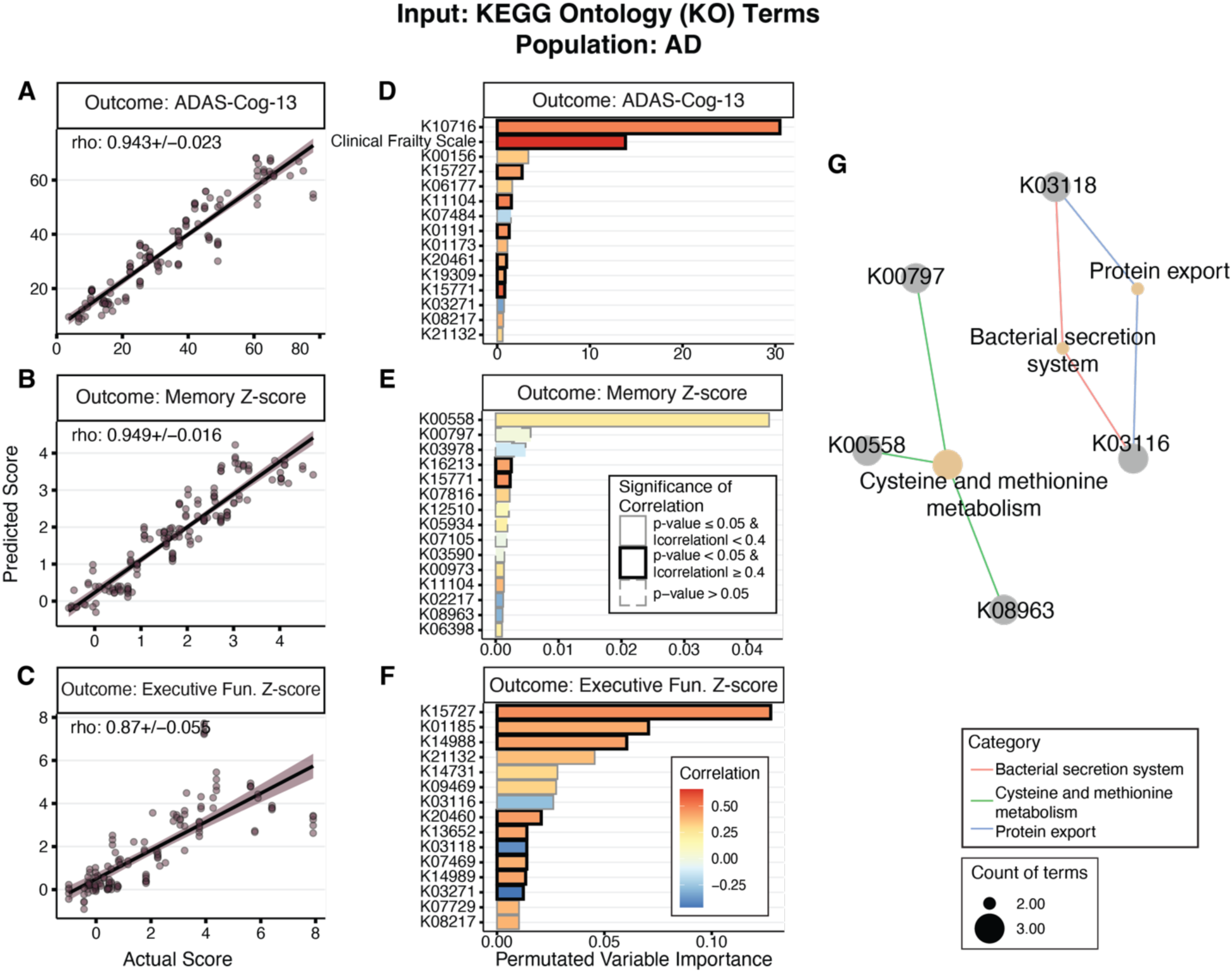
MERF regression models using KEGG Ontology (KO) term abundance and clinical covariates to predict cognitive outcomes in AD patient population. Actual versus predicted in AD population for models predicting ADAS-Cog-13 (A), memory z-score (B), and executive function z-score (C). Permutated variable importance for the final model for the ADAS-Cog-13 (D), memory z-score (E), and executive function z-score (F). Bar color corresponds to the correlation coefficient of each variable the indicated cognitive outcome. Significance of correlation is indicated by either a dashed (where p-value > 0.05) or a solid outline (p-value <= 0.5). Magnitude of correlation is indicated by color of the outline with gray indicating correlation below an absolute value of 0.4 and black indicating the correlation coefficient is great or equal to 0.4.

## DISCUSSION

Here, we demonstrate that the longitudinal microbiome profile of the GAINS subjects is broadly consistent across cognitive statuses and yet associated with cognitive performance in impaired patients. Within the GAINS cohort, diversity does not differ based on cognitive status. Age, clinical frailty scale, and overall CDR were the most frequent predictors of microbial abundance in the GAINS cohort. ML models were used to identify specific microbes, metabolic pathways, and gene groups of the gut microbiome that predict cognitive scores in AD subjects. Our analysis using species and encoded metabolic pathways identified features with known impacts on AD and immune-modulating roles as highly predictive of AD-specific cognitive function.

From the species abundance modeling, our analysis highlights a strong association between cognitive performance and the abundance of health-associated Clostridia IV and XIVa, such as members of *Blautia* genus and *Lachnospiraceae* family. Clostridia cluster IV and XIVa produce short-chain fatty acids (SCFAs), particularly butyrate, which has anti-inflammatory properties and supports the gut-brain axis^16^. These bacteria help maintain gut barrier integrity, reduce systemic inflammation, and modulate immune responses, protecting against cognitive decline^39,40^. Collectively, the function of these health-associated bacteria promotes brain health and enhances cognitive function. As taxonomy does not necessarily represent the functional ability of the microbiome, we next sought to examine the functional microbiome through metabolic pathway-based analysis.

Cysteine and methionine metabolism were enriched in the cognitive function-associated gene groups selected by MERF models, the metabolite components of which are highly studied in AD due in part to the potential use of homocysteine as a blood-based biomarker of AD^41–44^. Microbial-encoded pathway-based models also selected numerous pathways involved LPS and peptidoglycan (PG) synthesis as important in predicting cognitive outcomes in the AD population. The products of these pathways are known immunoreactive molecules that cause peripheral inflammation, which is known to influence neurocognitive function [see review by^45^]. This is the first longitudinal study to link cognitive function in AD subjects with microbial features and provide avenues beyond simple species-based associations.

### Inflammatory properties of the AD microbiome may drive worse cognitive outcomes

In our AD cohort, we also found that depletion of the common commensal bacteria *Phocaeicola dorei* was informative of worse cognitive performance in ADAS-Cog-13 (ranked 7^th^) and executive functioning z-score (ranked 6^th^). The lipid A component of *P. dorei* LPS is tetra- and penta-acylated compared to hexa-acylated lipid A found in microbes like *E. coli*^46^. The number of acyl groups has been previously found to be a strong detriment of immune activation with penta- and tetra-acylated lipid A showing reduced TLR4 response^47,48^. In two different studies, the administration of *P. dorei* to mice decreased colon LPS and further displayed inhibitor activity on host innate immune signaling^48,49^.

*Bacteroides thetaiotaomicron* was identified as important for ADAS-Cog-13 (rank 11^th^) and EF z-score (ranked 1^st^), both of which showed worse scores trending with increased abundance. Commonly, health-associated gram-negative *B. thetaiotaomicron is* decreased in individuals with obesity^50^ and to protect from diet-induced obesity in female mice^51^. Like *P.*dorei, the lipid A of some strains of *B. thetaiotaomicron* are penta-acylated and induces a weak TLR4 response^52^. Despite the similarity between our top predictors, *B. thetaiotaomicron* and *P. dorei* show differing associations with cognitive measures. While *B. thetaiotaomicron* may be commonly studied, very little is known about the genes involved in its LPS generation, which has led to the recent finding that *B. thetaiotaomicron* may not produce LPS but instead lipooligosaccharide (LOS), which is shorter and smaller than LPS as it is missing the O-antigen^53^.

While the gram-negative *P. dorei* is negatively correlated with worse cognitive function, pathways involved in peptidoglycan synthesis showed increased abundance correlating with worse cognitive function. Two UDP-pentapeptide biosynthesis pathways (PWY-7953 and PWY-6387) that feed into peptidoglycan biosynthesis and a peptidoglycan pathway (PWY-6385) were selected by both the ADAS-Cog-13 and memory z-score models (Figure 4B.14). In the executive function z-score model a precursor of either peptidoglycan or LPS showed an inverse correlation where lower precursor biosynthesis was significantly associated with worse EF Z-scores. LPS has been postulated to play an exacerbating role in AD pathophysiology.

The exterior of gram-negative bacteria in the human gut possesses large amounts of LPS and PG (endotoxin) that act as potent stimulators of the innate immune system when bacterial cells die or release small vesicles. However, the extent and duration of an inflammatory response are determined in a complex and often microbe-specific manner^54^. While CNS inflammation is a known component of AD pathology, the role of peripheral inflammation has been gaining traction in the field due to evidence of elevated endotoxin levels originating from peripheral infections or gut barrier dysfunction contributing to AD pathophysiology by elevating LPS levels in the blood and brain^55–58^. Evidence of endotoxin mediation of AD pathophysiology is extensive.

Significantly increased LPS in the serum and plasma of individuals with AD compared to healthy aged match controls has been identified by multiple research groups^59–64^. Further, Loffredo and authors identified that LPS levels positively correlated with serum zonulin levels, a measure of gut barrier permeability^60^. Marizzoni and authors also assessed gut permeability but did not find a significant correlation with serum LPS^61^. Specifically relating to cognition, evidence in healthy humans and mouse models indicates the onset of cognitive dysfunction by LPS administration^56^. In mice, multiple studies have demonstrated a causal relationship between LPS administration in the periphery and severity of cognitive impairment^65^. In our analysis, the identified association of endotoxin-related pathways and cognitive outcomes is specific to the AD population and not seen in the MCI population (data not shown here). Further, bacteria with low TLR4 activity showed an inverse correlation with cognitive scores, suggesting that an increase in potent TLR4 activators is accompanied by a decrease in known weaker potentiators.

Conversely, our models also identified a strong association between impaired cognitive performance and the decreased abundance of Clostridia cluster IV and XIVa members, specifically those of the *Blautia* genus and *Lachnospiraceae* family. Depletion of members of these clusters has been found in patients with inflammatory bowel disease and allergic diseases^66–70^. Interestingly, a human-derived consortia of Clostridia cluster IV and XIVa members have been extensively studied for its ability to induce immune tolerance in the gut, supporting the consortia’s anti-inflammatory abelites^66,67,71^. We postulate that the combined depletion of anti-inflammatory clostridia species and a shift to a higher endotoxicity of the gut microbiota are associated with poor cognitive function via the promotion or exacerbation of already established AD pathophysiology.

### Gut microbiota-derived methionine and B-vitamin pathways suggest increased production capability correlating with cognitive impairment

Our pathway-based analysis identified vitamin B12 and methionine-related pathways, such as *S*-adenosyl-L-methionine salvage (PWY-6151), as significantly correlated with cognitive outcomes. Methionine is an essential amino acid not synthesized by humans that when converted to *S-*adenosyl-methionine (SAM), is used by methyltransferases as a universal methyl group donor. Donation of the methyl group from SAM produces *S-*adenosyl-homocysteine (SAH), which is converted back to SAM via the *S*-adenosyl-L-methionine salvage pathway identified as the 5^th^ most important pathway for prediction of EF z-score and showed positive correlation with worsening score. In *E. coli,* this pathway can use either a cobalamin-independent or -dependent pathway (vitamin B12), the latter of which occurs 100-fold faster^72,73^. Together, our pathway-based analysis depicts an increased abundance of pathways utilizing methionine and those that support the use of methionine through salvaging necessary co-factors, suggesting an abundance of methionine in the gut that may correlate with cognitive impairment.

Gene family analysis identified an enrichment of cysteine and methionine biosynthesis genes in our KO-terms selected as important in predicting cognitive outcomes. Interestingly, results from both wild-type and AD-model mice suggest that increased dietary methionine induces a neurotoxic effect through increased APP deposition and a heightened inflammatory and oxidative stress response^41,42,44,74^. In humans, methionine and its related metabolites have shown significant promise as a blood-based biomarker for AD^42,44^. Increased contribution of methionine from the microbiome may thus serve a role in exacerbating AD-induced cognitive impairment.

## Conclusions

AD is a devastating, progressive disease impacting millions worldwide, with few effective therapies to slow down progression. Building upon the field’s established associations between AD and the microbiome, we demonstrate that the depletion of anti-inflammatory bacteria and enrichment of inflammatory metabolites is highly predictive of cognitive outcomes in our AD population. This work highlights the complexity of the microbiota-gut-brain axis and how the microbiome community makeup might play a role in exacerbating cognitive decline.

## ACKNOWLEDGEMENTS

We would like to thank the administration and staff from the Clinical Research Center here at UMass Medical Center and the Center for Clinical and Translational Sciences at UMass Chan Medical School for clinical facilities that supported the GAINS cohort. We would also like to thank the clinical research staff that supported this work including Samuel N. Odjidja, Protiva Dutta, Patrick M McGrath, Imane Samari, Lethycia Romeiro, Abigail Lopes, and Danielle Ferdinand.

## FUNDING

This study was designed and carried out at the University of Massachusetts Chan Medical School. JPH was supported by an Alzheimer’s Association Grant (2019-AARG-NTF-641955) and NIH grants from the National Institute on Aging (grant numbers: 2019-AARG-NTF-641955, R01AG067483-01). This prospective cohort study was approved by the institutional review board at the University of Massachusetts Chan Medical School (IRB docket H00021745). VB acknowledges support from the Bill & Melinda Gates Foundations and from the National Institute of Allergy and Infectious Diseases U01 AI172987.

## DATA AVAILABILITY

Data will be deposited upon article acceptance into publicly available repository and are also available from the PI upon request.

## Notes

### Competing Interest Statement

The authors have declared no competing interest.

