## Supplemental Tables and Figures for "Microbiome functional gene pathways predict cognitive performance in older adults with Alzheimer’s disease"

SUPPLEMENTAL DATA

Supplemental Table 1: Daily medications taken by GAINS cohort participants, ordered from most to least prevalent.

| Class | Specific | n | % |
|---|---|---|---|
| **Statins** | | 104 | 46.6 |
| **Acid reducing medications** | PPI | 53 | 23.8 |
| | H2 Blocker | 13 | 5.8 |
| **Antidepressants** | | 38 | 17.0 |
| | SSRIs | 38 | 17.0 |
| | Nerve pain medications | 21 | 9.4 |
| **Beta blockers** | | 34 | 15.2 |
| | Beta 1 selective agents | 33 | 14.8 |
| | Nonselective beta blocker agents | 5 | 2.2 |
| | Beta 1 and 3 selective agonist agents | 2 | 0.9 |
| | Beta 2 selective agents | 1 | 0.4 |
| **ACE Inhibitors** | | 34 | 15.2 |
| **Thyroid replacement hormones** | | 29 | 13.0 |
| **Seizure Medications** | | 26 | 11.7 |
| | GABA Analogs | 17 | 7.6 |
| | Sulfamate Substitute Monosaccharides | 4 | 1.8 |
| **Antiplatelet Medications** | | 24 | 10.8 |
| **NSAIDs** | | 20 | 9.0 |
| **Antihistamines** | | 20 | 9.0 |
| **Calcium channel blockers** | | 19 | 8.5 |
| | Dihydropyridine calcium channel blockers | 18 | 8.1 |
| | Calcium Channel Modulators | 17 | 7.6 |
| | Non-dihydropyridine calcium channel blockers | 1 | 0.4 |
| **Diuretics** | | 19 | 8.5 |
| | Thiazide diuretics | 15 | 6.7 |
| | Loop diuretics | 7 | 3.1 |
| | Potassium sparing diuretics | 2 | 0.9 |
| **Cholinesterase inhibitors** | | 18 | 8.1 |

| | | | |
|---|---|---|---|
| **Oral Medications for Diabetes** | | 18 | 8.1 |
| | Biguanides | 16 | 7.2 |
| | Meglitinides | 6 | 2.7 |
| | Sulfonylureas | 6 | 2.7 |
| | Sodium glucose transporter (SGLT-2) inhibitors | 4 | 1.8 |
| | Dipeptidyl peptidase-4 (DPP-4) inhibitors | 1 | 0.4 |
| | Thiazolidinediones | 1 | 0.4 |
| **Memory Medications** | | 18 | 8.1 |
| **Angiotensin-2 receptor blockers** | | 17 | 7.6 |
| **NMDA Receptor Blockers** | | 16 | 7.2 |
| **Benzodiazepines** | | 16 | 7.2 |
| **Anticoagulation** | | 12 | 5.4 |
| **Barbiturates** | | 11 | 4.9 |
| **Probiotics** | | 9 | 4.0 |
| | *Lactobacillus rhamnosus* | 2 | 0.9 |
| | *Lactobacillus acidophilus* (solo) | 1 | 0.4 |
| | *Lactobacillus acidophilus* (combo) | 1 | 0.4 |
| **Injectable Diabetes medications** | Glucagon-like peptide-1 receptor agonists (GLP-1RAs) | 8 | 3.6 |
| | Injectable Insulin Medication | 5 | 2.2 |
| **Atypical Antipsychotics** | | 5 | 2.2 |
| **Oral corticosteroids** | | 3 | 1.3 |
| **Immunosuppressive** | | 3 | 1.3 |
| **Vasodilators** | | 3 | 1.3 |
| **Carboxamides** | | 2 | 0.9 |
| **Sulfonamides** | | 2 | 0.9 |
| **Chemotherapy Medication** | | 2 | 0.9 |

Supplemental Table 2: Results of linear mixed-effect models for prediction of diversity in GAINS samples

| |
|---|
| *Alpha Diversity ~ Sex + Age+ Sample Day+ Education+ Abx Use + Status+ (1∣study id)* |

| Variable | Value | Std Error | DF | p-value |
|---|---|---|---|---|
| **(Intercept)** | 26.1052995 | 10.194995 | 436 | 0.01078497 |
| **Sex (male)** | 0.35107479 | 1.53567024 | 126 | 0.81954004 |
| **Age** | -0.1125021 | 0.10697484 | 126 | 0.29496432 |
| **Sample Day** | 0.00038433 | 0.00142376 | 436 | 0.78733167 |
| **Education (years)** | 0.11895518 | 0.26083751 | 126 | 0.64913844 |
| **Status (Linear contrast)** | -0.6677121 | 1.6691115 | 126 | 0.68980413 |
| **Status (Quadratic contrast)** | -1.9096678 | 1.81606289 | 126 | 0.2950219 |
| **Abx usage (past 6months)** | -4.3632227 | 1.76756727 | 126 | 0.0149088 |

Supplemental Table 3: All MERF regression model statistics presented as median [M.A.D].

| Outcome | Feature type | Population | Correlation Coef. | RMSE |
|---|---|---|---|---|
| ADAS-Cog-13 | Species Abundance | HC | 0.640 [0.05] | 3.028 [0.19] |
| | | MCI | 0.873 [0.04] | 3.574 [0.6] |
| | | AD | 0.941 [0.03] | 7.519 [1.1] |
| Memory Z-score | Species Abundance | HC | 0.626 [0.03] | 0.333 [0.01] |
| | | MCI | 0.825 [0.07] | 0.394 [0.05] |
| | | AD | 0.929 [0.01] | 0.505 [0.06] |
| Executive Function Z-score | Species Abundance | HC | 0.718 [0.02] | 0.399 [0.06] |
| | | MCI | 0.852 [0.04] | 0.446 [0.06] |
| | | AD | 0.846 [0.04] | 1.569 [0.1] |
| ADAS-Cog-13 | Metabolic Pathways | HC | 0.627 [0.06] | 2.995 [0.2] |
| | | MCI | 0.874 [0.04] | 3.508 [0.7] |
| | | AD | 0.940 [0.02] | 6.493 [1.9] |
| Memory Z-score | Metabolic Pathways | HC | 0.605 [0.02] | 0.325 [0.03] |
| | | MCI | 0.813 [0.07] | 0.388 [0.04] |
| | | AD | 0.944 [0.02] | 0.444 [0.08] |
| Executive Function Z-score | Metabolic Pathways | HC | 0.720 [0.02] | 0.400 [0.04] |
| | | MCI | 0.806 [0.04] | 0.483 [0.07] |
| | | AD | 0.882 [0.04] | 1.312 [0.2] |
| ADAS-Cog-13 | KO Terms | HC | 0.611 [0.04] | 3.105 [0.1] |
| | | MCI | 0.876 [0.03] | 3.323 [0.5] |
| | | AD | 0.935 [0.03] | 6.279 [1.5] |
| Memory Z-score | KO Terms | HC | 0.629 [0.03] | 0.322 [0.03] |
| | | MCI | 0.805 [0.06] | 0.394 [0.02] |

| | | | | |
|---|---|---|---|---|
| | | AD | 0.953 [0.01] | 0.474 [0.04] |
| Executive Function Z-score | KO Terms | HC | 0.714 [0.03] | 0.414 [0.04] |
| | | MCI | 0.795 [0.05] | 0.455 [0.04] |
| | | AD | 0.873 [0.04] | 1.461 [0.2] |

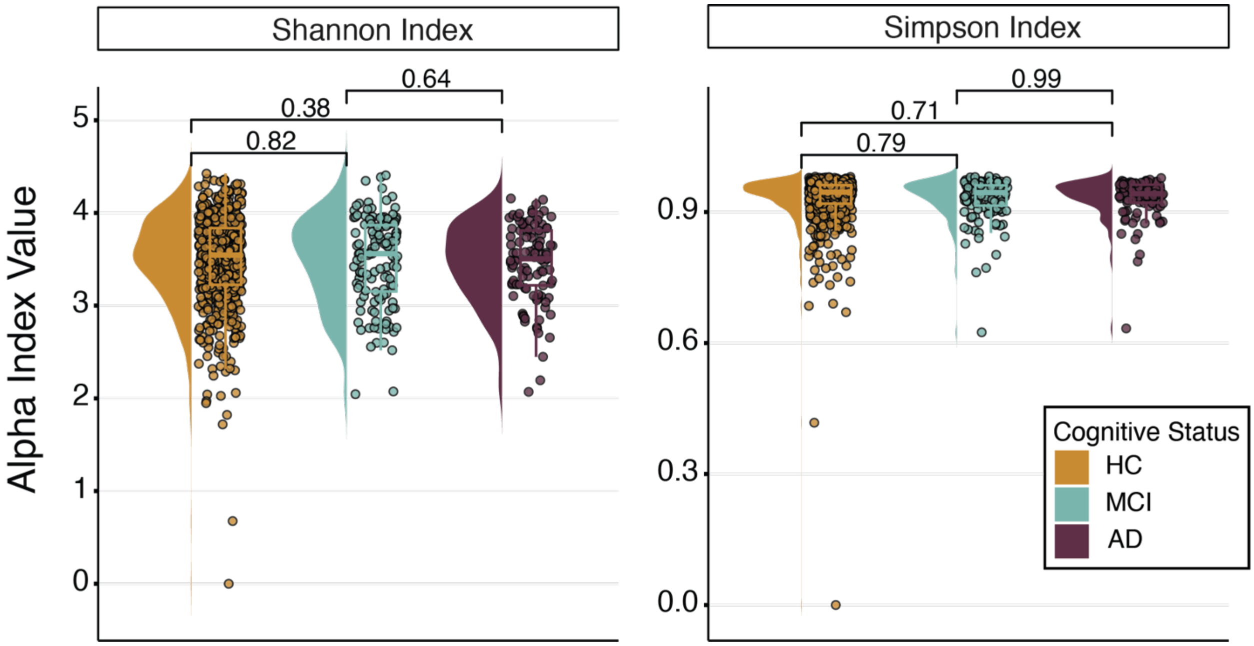


**Supplemental Figure 1: Alpha diversity indices.** Shannon and Simpson index values for GAINS participants across the three cognitive status sub populations; cognitively normal healthy controls (HC; orange yellow), mild cognitively impaired (MCI; teal), and diagnosed with Alzheimer's Disease (AD; violet).

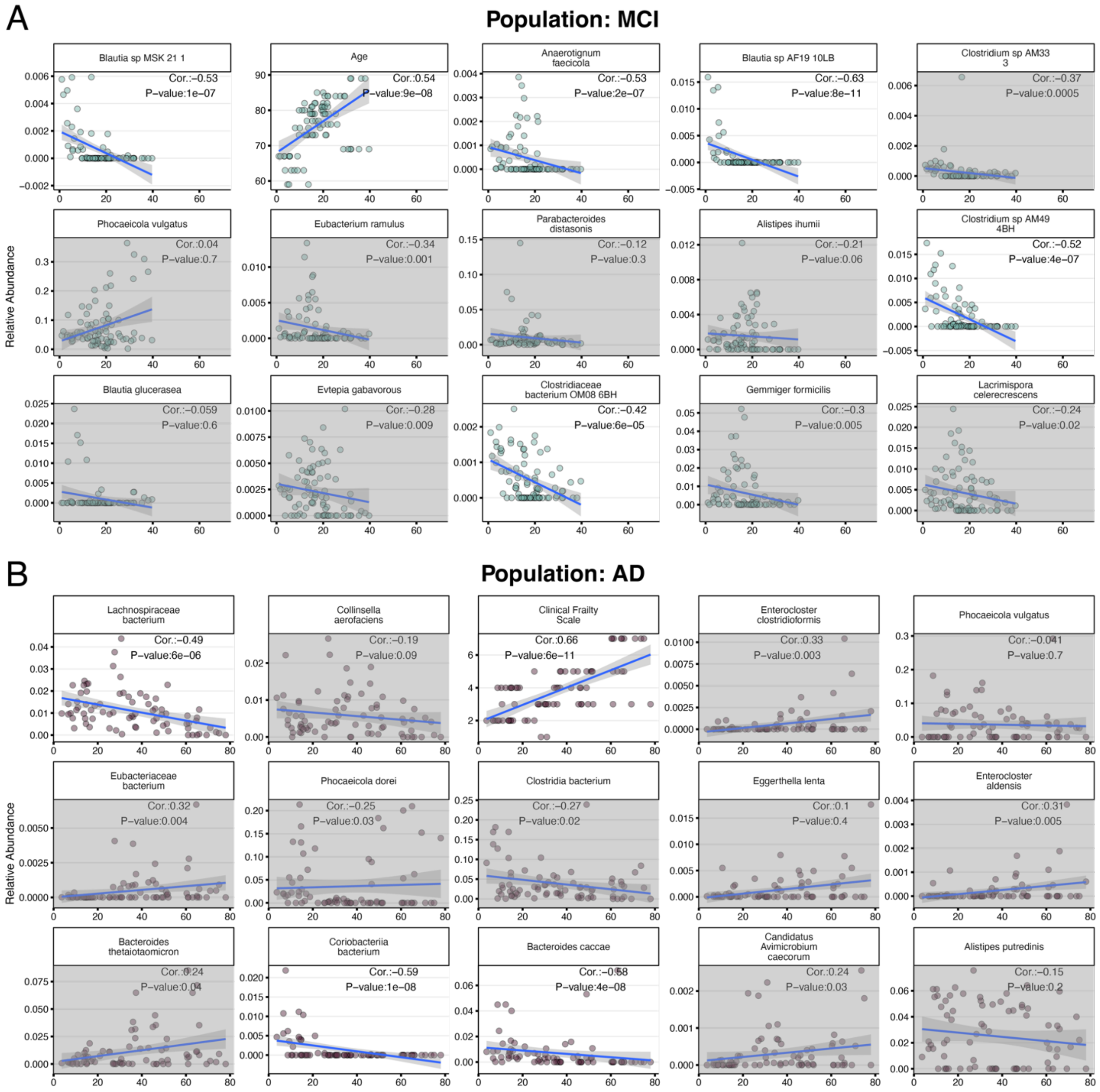


**Supplemental Figure 2: Top predictors selected by species abundance ADAS-Cog-13 MERF models.** Species abundance plotted against ADAS-Cog-13 scores for MCI population (A) and the AD population (B). Scores on x-axes. A grayed-out background indicates that the correlation of the variable with outcome was either not significant by p-value (p-value > 0.05) or that the absolute value of the correlation coefficient was less than 0.4.

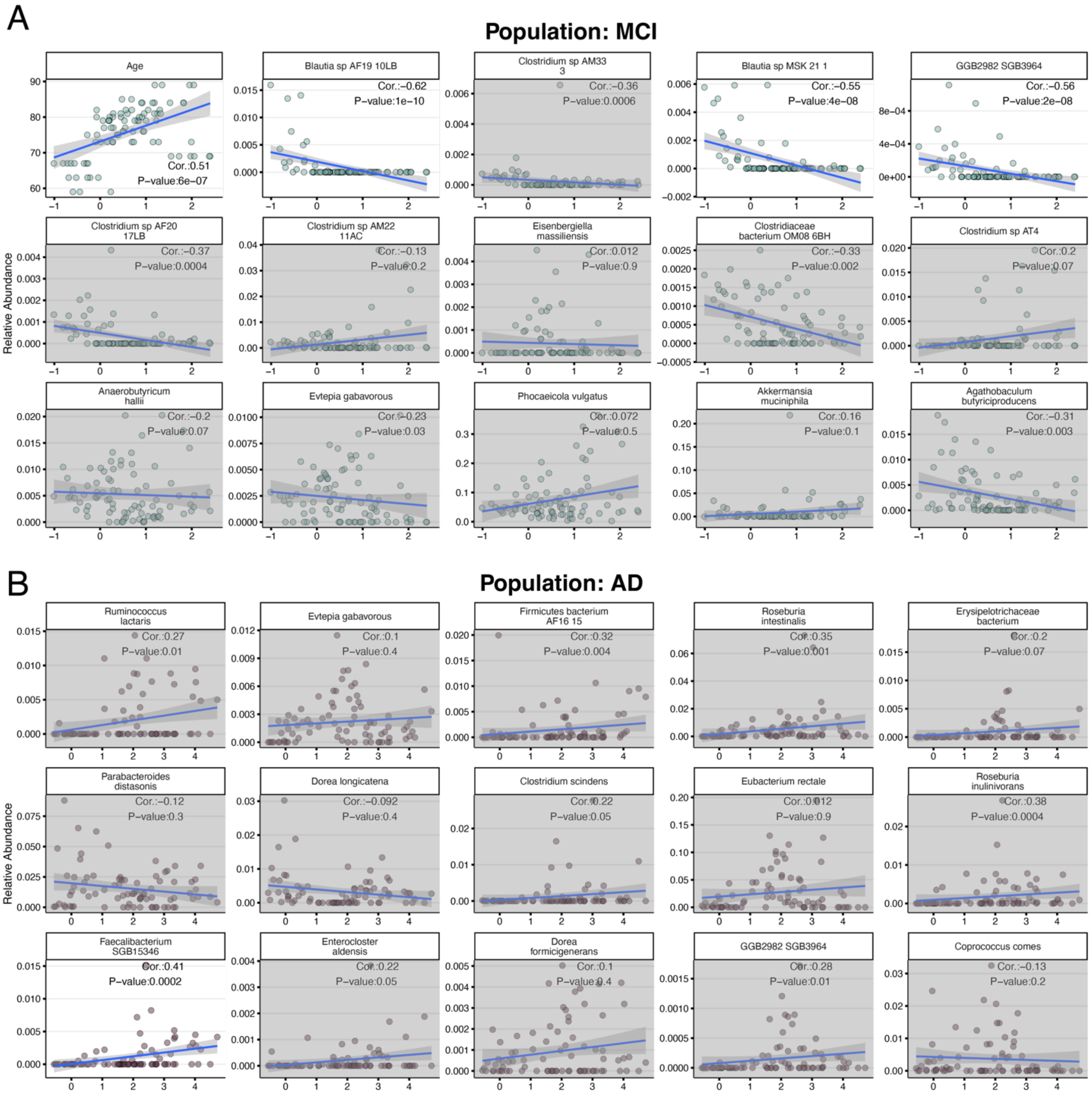


**Supplemental Figure 3: Top predictors selected by species abundance memory z-score MERF models.** Species abundance plotted against memory z-scores for MCI population (A) and the AD population (B). Scores on x-axes. A grayed-out background indicates that the correlation of the variable with outcome was either not significant by p-value (p-value > 0.05) or that the absolute value of the correlation coefficient was less than 0.4.

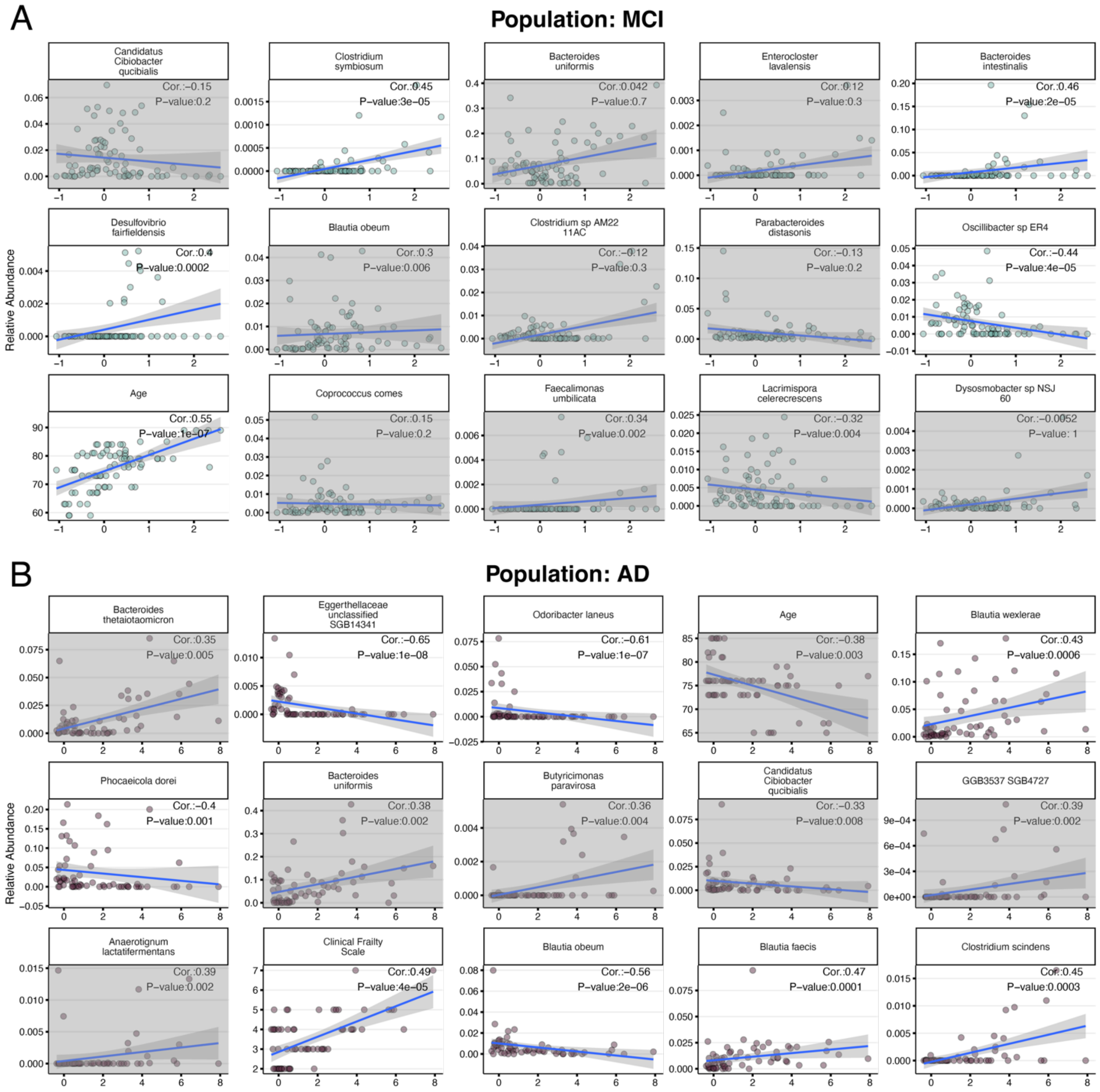


**Supplemental Figure 4: Top predictors selected by species abundance executive function z-score MERF models.** Species abundance plotted against EF Z-scores for MCI population (A) and the AD population (B). Scores on x-axes. A grayed-out background indicates that the correlation of the variable with outcome was either not significant by p-value (p-value > 0.05) or that the absolute value of the correlation coefficient was less than 0.4.

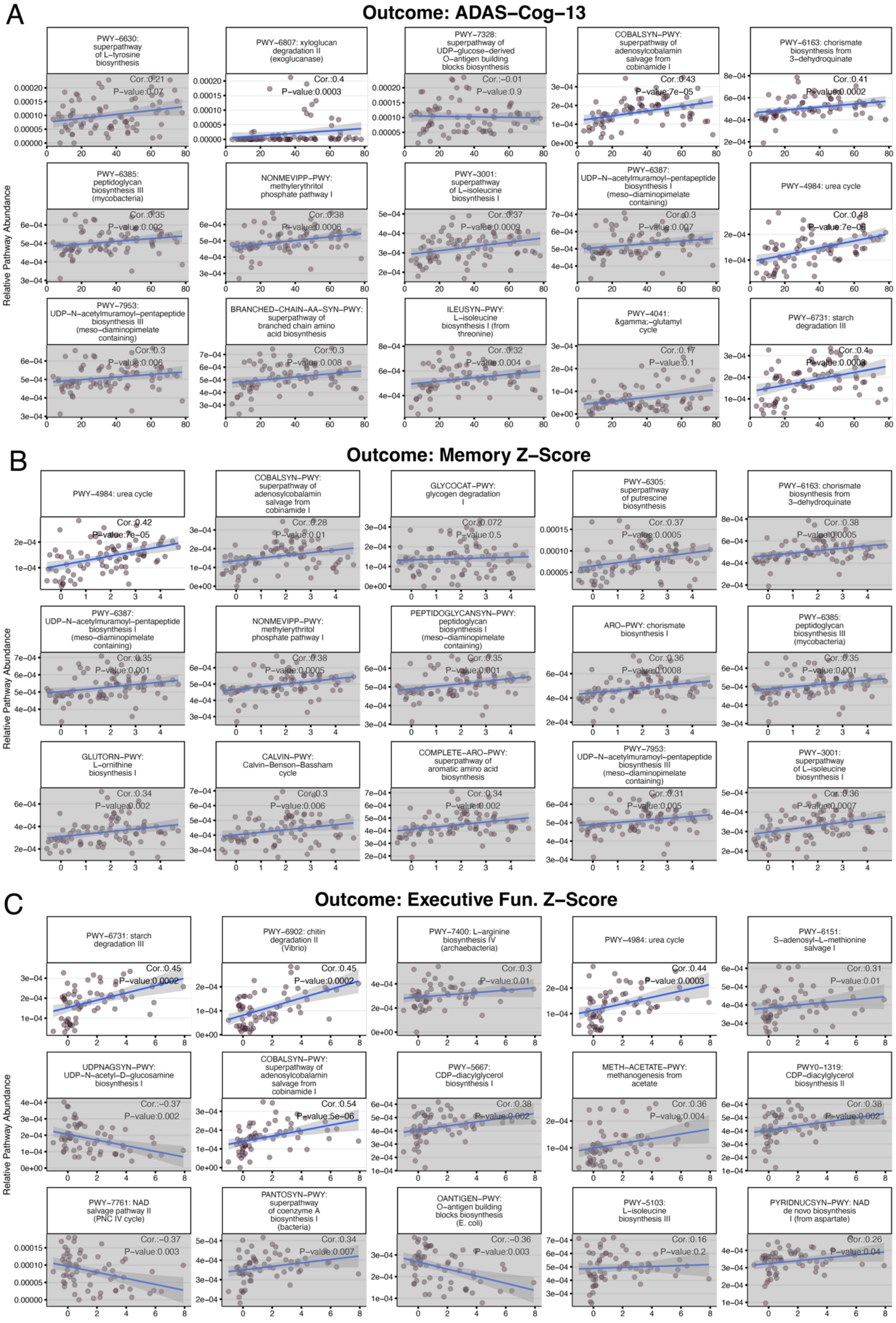
A
Outcome: ADAS−Cog−13
PWY−6630: superpathway of L−tyrosine biosynthesis
Cor.:0.21
P−value:0.07
PWY−6807: xyloglucan degradation II (exoglucanase)
Cor.:0.4
P−value:0.0003
PWY−7328: superpathway of UDP−glucose−derived O−antigen building blocks biosynthesis
Cor.:−0.01
P−value:0.9
COBALSYN−PWY: superpathway of adenosylcobalamin salvage from cobinamide I
Cor.:0.43
P−value:7e−05
PWY−6163: chorismate biosynthesis from 3−dehydroquinate
Cor.:0.41
P−value:0.0002
PWY−6385: peptidoglycan biosynthesis III (mycobacteria)
Cor.:0.35
P−value:0.002
NONMEVIPP−PWY: methylerythritol phosphate pathway I
Cor.:0.38
P−value:0.0006
PWY−3001: superpathway of L−isoleucine biosynthesis I
Cor.:0.37
P−value:0.0009
PWY−6387: UDP−N−acetylmuramoyl−pentapeptide biosynthesis I (meso−diaminopimelate containing)
Cor.:0.3
P−value:0.007
PWY−4984: urea cycle
Cor.:0.48
P−value:7e−06
PWY−7953: UDP−N−acetylmuramoyl−pentapeptide biosynthesis III (meso−diaminopimelate containing)
Cor.:0.3
P−value:0.006
BRANCHED−CHAIN−AA−SYN−PWY: superpathway of branched chain amino acid biosynthesis
Cor.:0.3
P−value:0.008
ILEUSYN−PWY: L−isoleucine biosynthesis I (from threonine)
Cor.:0.32
P−value:0.004
PWY−4041: γ−glutamyl cycle
Cor.:0.17
P−value:0.1
PWY−6731: starch degradation III
Cor.:0.4
P−value:0.0003
Relative Pathway Abundance
B
Outcome: Memory Z−Score
PWY−4984: urea cycle
Cor.:0.42
P−value:7e−05
COBALSYN−PWY: superpathway of adenosylcobalamin salvage from cobinamide I
Cor.:0.28
P−value:0.01
GLYCOCAT−PWY: glycogen degradation I
Cor.:0.072
P−value:0.5
PWY−6305: superpathway of putrescine biosynthesis
Cor.:0.37
P−value:0.0005
PWY−6163: chorismate biosynthesis from 3−dehydroquinate
Cor.:0.38
P−value:0.0005
PWY−6387: UDP−N−acetylmuramoyl−pentapeptide biosynthesis I (meso−diaminopimelate containing)
Cor.:0.35
P−value:0.001
NONMEVIPP−PWY: methylerythritol phosphate pathway I
Cor.:0.38
P−value:0.0005
PEPTIDOGLYCANSYN−PWY: peptidoglycan biosynthesis I (meso−diaminopimelate containing)
Cor.:0.35
P−value:0.001
ARO−PWY: chorismate biosynthesis I
Cor.:0.36
P−value:0.0008
PWY−6385: peptidoglycan biosynthesis III (mycobacteria)
Cor.:0.35
P−value:0.001
GLUTORN−PWY: L−ornithine biosynthesis I
Cor.:0.34
P−value:0.002
CALVIN−PWY: Calvin−Benson−Bassham cycle
Cor.:0.3
P−value:0.006
COMPLETE−ARO−PWY: superpathway of aromatic amino acid biosynthesis
Cor.:0.34
P−value:0.002
PWY−7953: UDP−N−acetylmuramoyl−pentapeptide biosynthesis III (meso−diaminopimelate containing)
Cor.:0.31
P−value:0.005
PWY−3001: superpathway of L−isoleucine biosynthesis I
Cor.:0.36
P−value:0.0007
Relative Pathway Abundance
C
Outcome: Executive Fun. Z−Score
PWY−6731: starch degradation III
Cor.:0.45
P−value:0.0002
PWY−6902: chitin degradation II (Vibrio)
Cor.:0.45
P−value:0.0002
PWY−7400: L−arginine biosynthesis IV (archaebacteria)
Cor.:0.3
P−value:0.01
PWY−4984: urea cycle
Cor.:0.44
P−value:0.0003
PWY−6151: S−adenosyl−L−methionine salvage I
Cor.:0.31
P−value:0.01
UDPNAGSYN−PWY: UDP−N−acetyl−D−glucosamine biosynthesis I
Cor.:−0.37
P−value:0.002
COBALSYN−PWY: superpathway of adenosylcobalamin salvage from cobinamide I
Cor.:0.54
P−value:5e−06
PWY−5667: CDP−diacylglycerol biosynthesis I
Cor.:0.38
P−value:0.002
METH−ACETATE−PWY: methanogenesis from acetate
Cor.:0.36
P−value:0.004
PWY0−1319: CDP−diacylglycerol biosynthesis II
Cor.:0.38
P−value:0.002
PWY−7761: NAD salvage pathway II (PNC IV cycle)
Cor.:−0.37
P−value:0.003
PANTOSYN−PWY: superpathway of coenzyme A biosynthesis I (bacteria)
Cor.:0.34
P−value:0.007
OANTIGEN−PWY: O−antigen building blocks biosynthesis (E. coli)
Cor.:−0.36
P−value:0.003
PWY−5103: L−isoleucine biosynthesis III
Cor.:0.16
P−value:0.2
PYRIDNUCSYN−PWY: NAD de novo biosynthesis I (from aspartate)
Cor.:0.26
P−value:0.04
Relative Pathway Abundance

**Supplemental Figure 5: Top predictors selected by pathway abundance models for each outcome in the AD population.** Selected pathway abundance in the AD patient population versus ADAS-Cog-13 (A), memory z-score (B), and executive function z-score (C). Scores on x-axes. A grayed-out background indicates that the correlation of the variable with outcome was either not significant by p-value (p-value > 0.05) or that the absolute value of the correlation coefficient was less than 0.4.
